# Octave plots for visualizing diversity of microbial OTUs

**DOI:** 10.1101/389833

**Authors:** Robert C. Edgar, Henrik Flyvbjerg

## Abstract

Next-generation sequencing of marker genes such as 16S ribosomal RNA is widely used to survey microbial communities. The abundance distribution (AD) of Operational Taxonomic Units (OTUs) in a sample is typically summarized by alpha diversity metrics, e.g. richness and entropy, discarding information about the AD shape. In this work, we describe octave plots, histograms which visualize the shape of microbial ADs by binning on a logarithmic scale with base 2. Optionally, histogram bars are colored to indicate possible spurious OTUs due to sequence error and cross-talk. Octave plots enable assessment of (a) the shape and completeness of the distribution, (b) the effects of noise on measured diversity, (c) whether low-abundance OTUs should be discarded, (d) whether alpha diversity metrics and estimators are reliable, and (e) the additional sampling effort (i.e., read depth) required to obtain a complete census of the community. The utility of octave plots is illustrated in a re-analysis of a prostate cancer study showing that the reported core microbiome is most likely an artifact of experimental error.

## Introduction

Metagenomics by next-generation sequencing of marker genes such as 16S ribosomal RNA (rRNA) has revolutionized the study of microbial communities in environments ranging from the human body (Cho and Blaser, 2012; Pflughoeft and Versalovic, 2012) to oceans (Moran, 2015) and soils (Hartmann et al., 2014). Data analysis in such studies typically assigns reads to clusters of similar sequences called Operational Taxonomic Units (OTUs). Alpha diversity, i.e. diversity of a single sample or community, is often characterized by a number (*metric*) calculated from the set of OTU abundances. For example, richness is the number of OTUs, and Shannon entropy (Shannon, 1948) is a function of the OTU frequencies. More detailed views of abundance distributions (ADs) are provided by AD plots (Preston, 1948; Whittaker, 1965), which are routinely used to show species diversity in traditional biodiversity studies (Magurran, 2004) but are almost entirely absent from the metagenomics literature.

While experimental error can largely be neglected in traditional studies, metagenomic OTUs are often spurious due to sequence errors (Edgar, 2013; Huse et al., 2010) and cross-talk (Carlsen et al., 2012; Edgar, 2017a), which can lead to grossly incorrect estimates of diversity (Edgar, 2017b). Biases favoring or disfavoring observations of some groups are recognized in traditional biodiversity (for example, larger species are easier to see), but are often considered to be inconsequential in communities of similar organisms (e.g., birds). In marker gene metagenomics, biases due to mismatches with PCR primers and variations in gene copy number are more severe, causing abundances of gene sequences in the reads to have low correlation with species abundances (Edgar, 2017c).

In both traditional and metagenomic studies, some groups (species or OTUs) may be missing because of insufficient observations; we will refer to this as *incomplete enumeration* to avoid over-use of the terms sample and sampling. Reviewing an AD plot enables an assessment of the shape of an AD and whether it is complete or incomplete. This assessment is informed by comparison with shapes generated by mathematical models. Many such models have been proposed (Magurran, 2004), including the log-series (Fisher et al., 1943) and log-normal distribution (Preston, 1948). The log-normal model is based on a probability density function for abundances of all groups, including both observed and unobserved. By contrast, Fisher’s log-series is a formula which is designed to model a substantially *incomplete* enumeration, i.e. one where many groups are unobserved (Fisher et al., 1943). An incomplete enumeration of a log-normal AD increasingly resembles a log-series as the number of observations decreases (Fig. 1). Log-normal is attractive empirically and theoretically because most observed macro-ecology ADs are consistent with an approximately log-normal community AD, and by the central limit theorem a community’s AD will be log-normal if its abundances are determined by many independent random factors (May, 1975). In our view, the log-normal is a more general and therefore superior model because a log-series is not designed to model complete enumerations and is well-approximated by an incomplete log-normal, while log-normal can model both complete and incomplete enumerations and an incomplete log-normal cannot be well-approximated by a log-series.

**Fig. 1.**
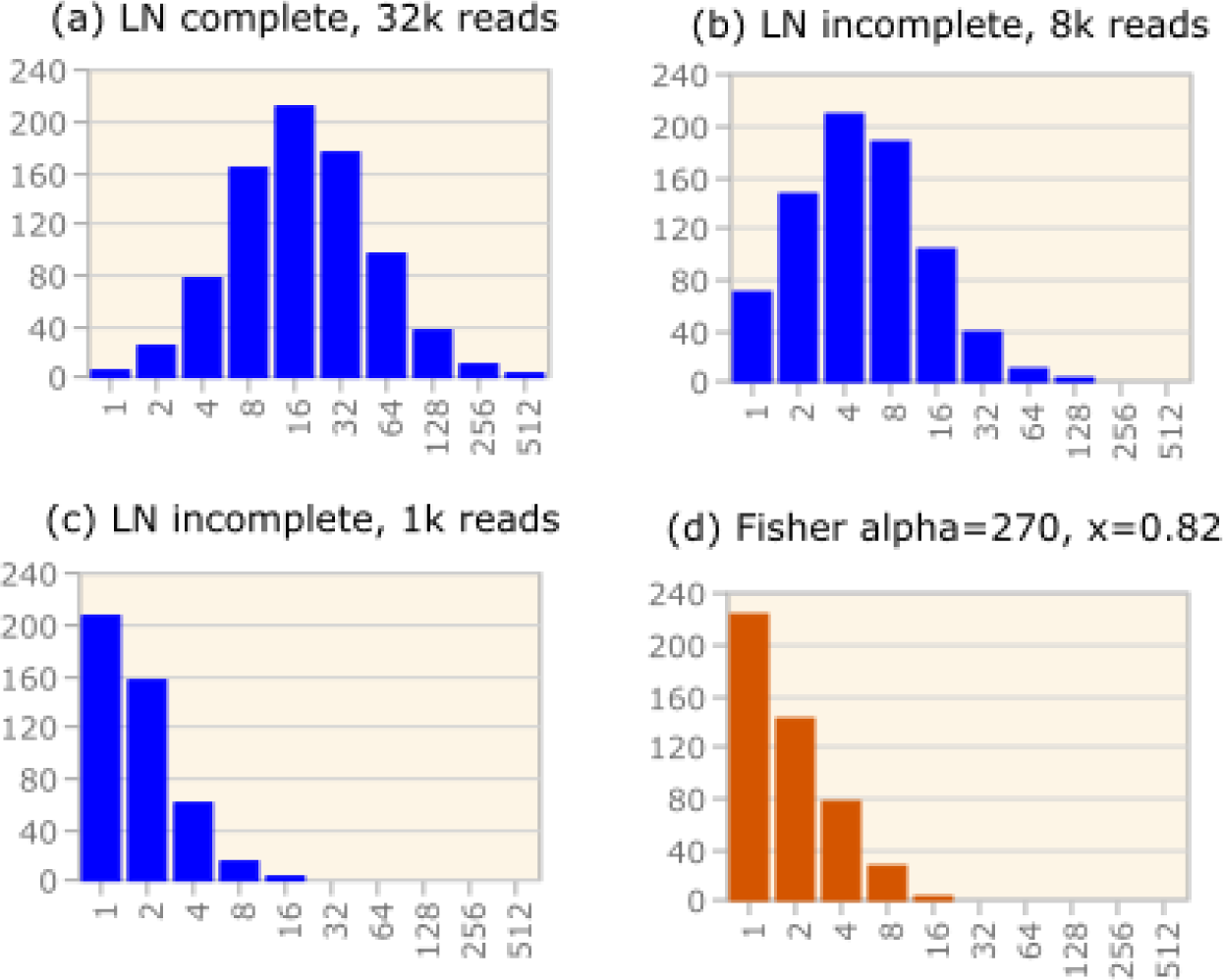
Simulated log-normal (blue) and Fisher log-series (brown) distributions. Panel (a) shows a complete enumeration of a log-normal distribution generated with parameters *μ*=4.6, *σ*=1.5, *G*=*G_obs_*=32k. Panels (b) and (c) show incomplete distributions obtained from the distribution in (a) by taking subsets of the observations (reads) of sizes *G_obs_*=8k and 1k respectively. Panel (d) is a log-series with parameters *α*=270, *x*=0.82. Note that (c) and (d) are similar, illustrating that a log-series resembles an incomplete log-normal where the peak is not apparent.

In this work, we introduce octave plots for visualizing abundance distributions of OTUs obtained by next-generating sequencing. Octave plots are modified versions of Preston plots with bin boundaries chosen to optimize preservation of the distribution shape and optional coloring to indicate possible spurious OTUs.

## Methods

### Histogram binning

By our definition, an octave plot is a histogram where the height of the *k*th bar (*k* = 0, 1…) is proportional to the number of OTUs with an abundance (*r*) in the range *r*=2^*k*^, 2^*k*^+1 … 2^*k*+1^−1. Note that two different numbers double from one bin to the next: (a) the minimum abundance, and (b) the range of abundances in the bin. Thus, on a logarithmic scale, bins are uniformly spaced and have the same width, which optimizes shape preservation with incomplete enumerations. Consider for example the effect of discarding a randomly-chosen half of the reads. To a first approximation, the abundance of every OTU will be halved and this has the effect of shifting all counts in the histogram, and hence all bars, one bin to the left. Discarding all but ½^*m*^ of the reads is roughly equivalent to moving bars *k*≥*m* to the left by *m* bins, preserving their shape, while bars *k*<*m* are moved past the *y* axis and disappear. Our definition of the bin boundaries has previously been considered, e.g. by (Gray et al., 2006), but is almost never used in practice. Plots described by Preston (Preston, 1948), a popular textbook (Magurran, 2004), and a claimed improvement on Preston’s method (Williamson and Gaston, 2005) all have bins with irregular sizes and spacing on a logarithmic scale. For example, Magurran’s first and second bins both contain exactly two abundances (*r*=1 and *r*=2 in the first bin, *r*=3 and *r*=4 in the second bin) while subsequent bins double in size. The first two bins proposed by (Williamson and Gaston, 2005) both contain exactly one abundance (*r*=1 in the first bin, *r*=2 in the second) while the third bin contains three abundances (*r*=3…5). Irregular bins can cause misleading distortions of shape such as peaks which do not correspond to the mean of the underlying log-normal distribution (Fig. 2).

**Fig. 2.**
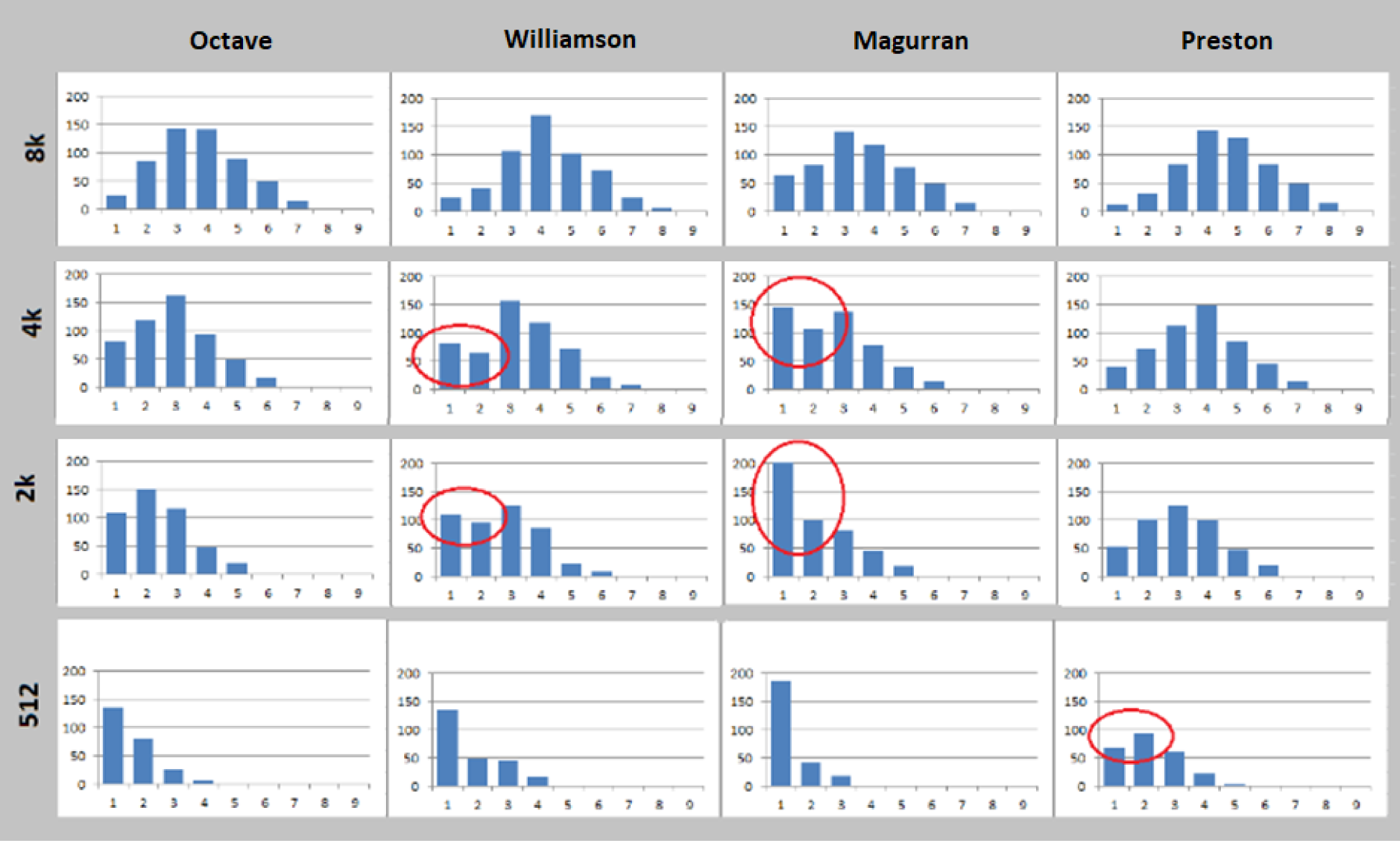
Shape distortions arising from different binning methods. All histograms in the figure were generated from the same simulated log-normal abundance distribution. The histograms in a given row are generated from an identical incomplete enumeration of this distribution. From top to bottom, the enumerations contain 8k, 4k, 2k and 512 reads, respectively. Binning rules are: *Octave* (this paper), *Williamson* (Williamson and Gaston, 2005), *Magurran* (Magurran, 2004) and *Preston* (Preston, 1948). Shape distortions are circled in red. Octave binning is the only method that preserves the shape without misleading distortions. Bins are labeled 1, 2 … 9 in order of increasing abundance.

With 16S rRNA OTUs, binning mitigates the problem that read abundance correlates poorly with species abundance. The gene count observed in most species ranges from one to nine per genome with roughly uniform frequencies (Edgar, 2017c). If three is chosen as the canonical standard, the abundance correction factor for gene count therefore ranges from 1/3 to 3, but cannot be usefully predicted from OTU sequences due to the sparse coverage of currently available reference databases (Edgar, 2017c). From this perspective, it would be preferable to use a logarithm base >2 as this would be more likely to assign an OTU to its correct bin. However, ADs are compressed into fewer histogram bars when using base 3 or higher, which we found to give less recognizable shapes for distributions encountered in practice. We therefore chose to use base 2 by default.

### Spurious OTUs due to sequence error

Reads with incorrect sequences can inflate diversity. For example, if the conventional 97% threshold is used, then a sequence with >3% errors will induce a spurious OTU. The correct sequence is likely to have much higher abundance because reads with many errors are rare. Thus, if a low-abundance OTU (*L*) has high sequence similarity with a high-abundance OTU (*D*), this suggests that *L* could be a spurious OTU for which *D* is the correct sequence. Let *Ab*(*x*) be the abundance of OTU *x*, and *Pctid*(*x*, *y*) be the sequence identity of OTUs *x* and *y* expressed as a percentage. If there is an OTU *D* such that *Ab*(*D*)/*Ab*(*L*) is large and *Pctid* (*D*, *L*) is high, then *L* is potentially spurious. We quantified this by classifying an OTU *L* as *strongly noisy* (i.e., there is strong evidence that *L* is spurious) if there is an OTU *D* such that (*Ab*(*D*)/*Ab*(*L*) ≥ *W_strong_ and Pctid*(*D*, *L*) ≥ *D_strong_*), and otherwise *weakly noisy* (i.e., weak evidence that *L* is spurious) if there is a *D* such that (*Ab*(*D*)/*Ab*(*L*) ≥ *W_weak_ and Pctid* (*D*, *L*) ≥ *D_weak_*), where by default *W_weak_*=32, *W_strong_*=256, *D_weak_*=92% and *D_strong_*=95%. Optionally, an octave plot bar is colored to indicate the fraction of OTUs in its bin which are weakly noisy (orange) or strongly noisy (red).

### Upper bound on diversity

Low-abundance read sequences, especially singletons, are enriched for errors which induce spurious OTUs and are therefore discarded by algorithms which aim to report accurate OTU sequences (Callahan et al., 2016; Edgar, 2013, 2017d). This strategy may underestimate the number of valid low-abundance OTUs, causing distortion of the reported AD. Pooling reads from all samples can mitigate this issue (Edgar, 2017d), for example by enabling detection of a singleton OTU in a given sample providing that it is present in sufficiently many other samples. However, some distortion may remain after pooling. An upper bound on the correct number of OTUs in each bin can be obtained by keeping all read sequences that pass a quality filter. The *surplus* of an octave plot bin is the increase in the number of OTUs when all read sequences are included, which gives an upper bound on the correct value. Optionally, the surplus is indicated by a gray segment at the top of an octave plot bar.

### Spurious abundances due to cross-talk

Reads are often assigned to the wrong sample due to cross-talk, which can cause spurious abundances, i.e. counts which should be zero (Carlsen et al., 2012; Edgar, 2018). Optionally, an octave plot bar is colored to indicate the fraction of OTUs in its bin which have UNCROSS2 scores (Edgar, 2018) that are ≥0.1 (weak cross-talk, yellow) or ≥0.4 (strong cross-talk, pink).

### Simulation of log-normal and log-series

We simulated a log-normal abundance distribution using a Gaussian probability density function (PDF) specified by parameters *μ* (mean) and *σ* (standard deviation). The size of the population was specified by *S*, the total number of species, including those not observed, or *M*, its total number of individuals. A simulated observed distribution was obtained by specifying a number of observations (reads) (*N*). A log-series was simulated using a distribution defined by parameters *α* and *x* where the expected number of OTUs with observed abundance *r* is *α x*^*r*^/*r*.

### Representative studies

To investigate representative OADs, we used reads of samples from the human vagina (Virtanen et al., 2017), a phytoplankton bloom (Parulekar et al., 2017) and soil (Carini et al., 2017). We generated OTU tables using the current recommended UPARSE (Edgar, 2013) protocol (https://drive5.com/usearch/manual/uparse_pipeline.html, accessed 1st August 2018), which discards singleton read sequences, i.e. sequences observed exactly once per dataset, and (2) the same protocol as (1) except that singletons reads are retained (*UPARSE+1*). Octave plots were created using the *otutab_octave* command in USEARCH (Edgar, 2010) version 11.0.

### Prostate cancer microbiome

We used reads of the 16S rRNA V4 hypervariable region from the prostate cancer microbiome study reported in (Yow et al., 2017). A minimum OTU abundance threshold was determined by reviewing octave plots (see Results), which was implemented using the *otutab_trim* command in USEARCH (Edgar, 2010) version 11.0, and a putative core microbiome was identified using the *otutab_core* command.

### QIIME open-reference OTUs

This recommended OTU protocol for QIIME (Caporaso et al., 2010) v1.9 is open-reference clustering (Rideout et al., 2014), which has been shown to generate large numbers of spurious OTUs on mock community tests (Edgar, 2017e). To investigate the ADs generated by open-reference clustering from *in vivo* datasets, we generated octave plots from OTUs generated by the QIIME v1.9 *pick_open_reference_otus.py* script after quality filtering by *split_libraries_fastq.py*.

### Mock community test

For testing on an artificial (*mock*) community of known composition, we used 16S rRNA reads of the Staggered1 sample of (Bokulich et al., 2013), which contains 21 strains with designed abundances ranging over three orders of magnitude.

## Results

### Simulated distributions

Some representative simulations are shown in Fig. 1. Panel (a) shows a complete log-normal distribution generated with parameters *μ*=4.6, *σ*=1.5, *M*=*N*=64k. Panels (b) and (c) show incomplete distributions obtained from the distribution in (a) with *N* (number of reads) 8k and 1k respectively. To a good approximation, the effect of reducing the number of reads is to move the histogram leftwards but otherwise preserve its shape. From (a) to (b) the shape is moved leftwards by log_2_(32k/8k)=2 bins, and from (b) to (c) it is moved by log_2_(8k/1k)=3 bins. In (b), the peak of the bell curve is still visible, but in (c) the peak has moved to bin 1 so that the histogram is monotonically decreasing. When the peak is at bin 1 or has moved to the left of the *y* axis, a log-normal distribution resembles a Fisher log-series, as illustrated by panel (d) which shows a log-series with parameters *α*=270, *x*=0.82.

### Shapes observed in practice

We found that ADs of UPARSE OTUs from *in vivo* datasets could be divided into four categories which we denote *C*, *T*, *J* and *A*, with representative examples shown in Fig. 3. *C* (“complete”) resembles a bell curve which we interpret as implying that the true distribution is well-modeled by a complete log-normal. *T* (‘truncated’) resembles a truncated bell curve where a roughly symmetrical peak is visible; we interpret this as an incompletely enumerated log-normal. *J* is a decreasing distribution which lacks a peak, resembling the letter “J” rotated 90° clockwise or a hockey stick. An AD of type *J*, which in our experience is the most common type, can be well-modeled by a log-series or a truncated log-normal. ADs of type *A* (“anomalous”) have irregular shapes. One possible biological explanation for an AD of type *A* is that the sample contains multiple independent niches or communities with minimal interaction.

**Fig. 3.**
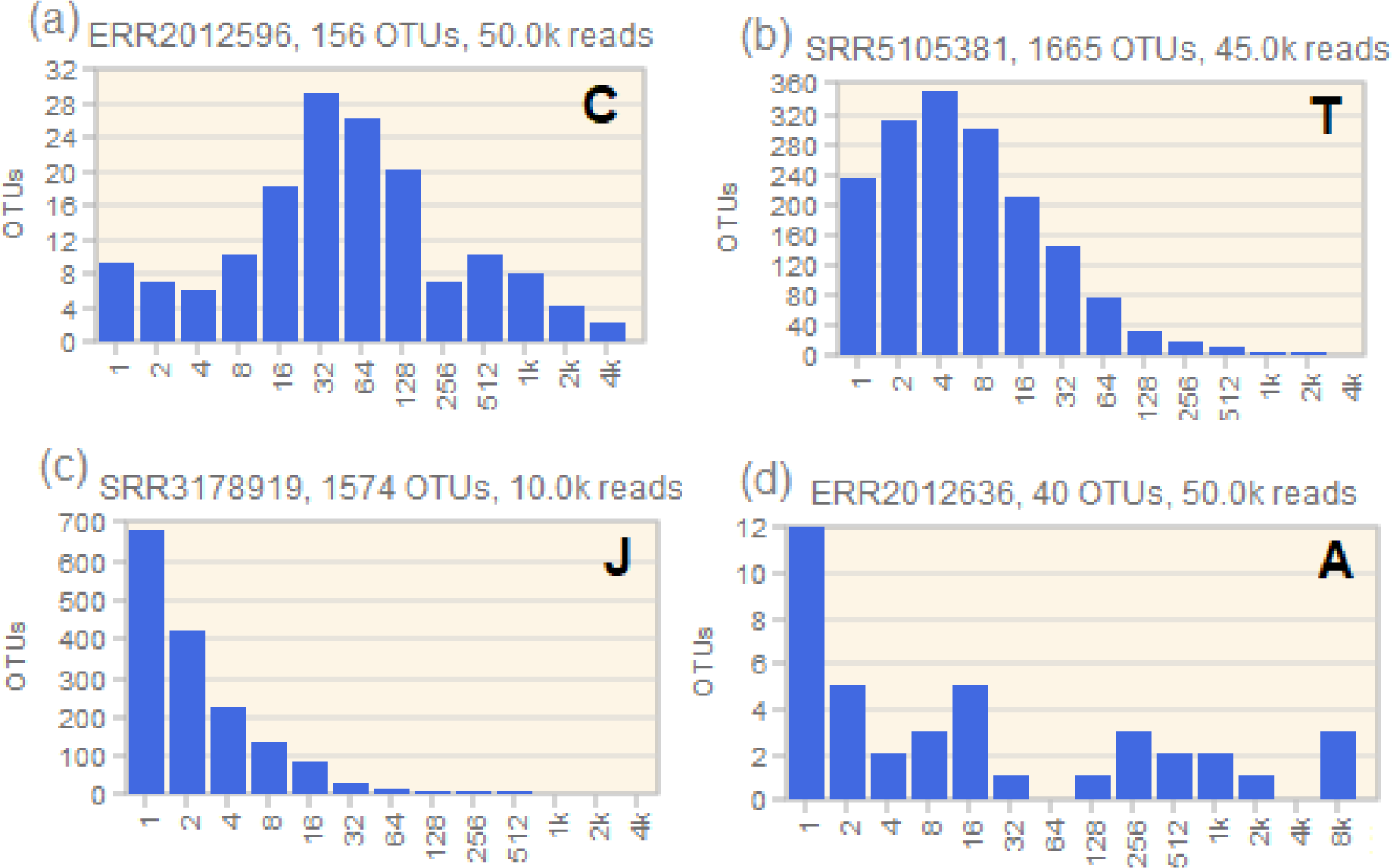
Examples of distributions observed in practice. Octave plots generated from UPARSE OTUs. (a) Vagina sample ERR2012596 from (Virtanen et al., 2017), (b) phytoplankton bloom sample SRR5105381 from (Parulekar et al., 2017), (c) soil sample SRR3178919 from (Carini et al., 2017) and (d) vagina sample ERR2012636 from (Virtanen et al., 2017). These distributions are classified as *C* (complete), *T* (truncated), *J* (hockey stick or *J*-shaped), and *A* (anomalous), respectively.

### Putative prostate cancer core microbiome

We use the suffix “k” to indicate multiples of 2^10^ = 1,024 and label a histogram bar by its minimum abundance, so e.g. bar 16 shows the number of OTUs with abundances in the range 16 … 31. The octave plot for a representative sample from (Yow et al., 2017) is shown in Fig. 4, which we interpret as follows. Bars 128 through 4k exhibit a bell curve shape consistent with an approximately log-normal interacting ecosystem. One very high-abundance OTU (75,902 reads = 61% of the sample) is an outlier which was identified by SINTAX (Edgar, 2016) as *Escherichia/Shigella* with bootstrap confidence 0.94. As shown by the coloring in Fig. 4, most OTUs in bins 1 through 64 (i.e., abundances 1 … 127) were identified as potentially spurious due to sequence error or cross-talk. We therefore chose 100 reads per sample as the minimum abundance required to confidently identify an OTU as present, which is substantially higher than the UPARSE defaults (minimum two reads per dataset and one read per sample). We believe that a high rate of spurious low-abundance OTUs is plausible for this dataset due to low biomass and deep sequencing (125k reads/sample after rarefaction). This analysis underscores that default parameters for an analysis pipeline may be sub-optimal for any given dataset. Using UPARSE+1, which maximizes sensitivity by retaining singleton read sequences at the possible expense of reporting an increased number of spurious OTUs, we found that *Escherichia/Shigella* was the only OTU with ≥100 reads in at least 95% of samples. Two other OTUs were present in all samples and an additional three OTUs were present in 95% of samples. The maximum abundances of these five OTUs ranges from 44 to 11,975 while the minimum non-zero abundance is one in all cases, which is consistent with cross-talk. The presence of these five OTUs therefore cannot be reliably established in samples where their abundances are low. Thus, six is an upper bound on the number of OTUs found in at least 95% of samples, of which five are likely to be false positives (i.e., in fact are present in <95% of samples). In conclusion, the data does not adequately support the claim made by (Yow et al., 2017) that 18 OTUs are present in at least 95% of samples.

**Fig. 4.**
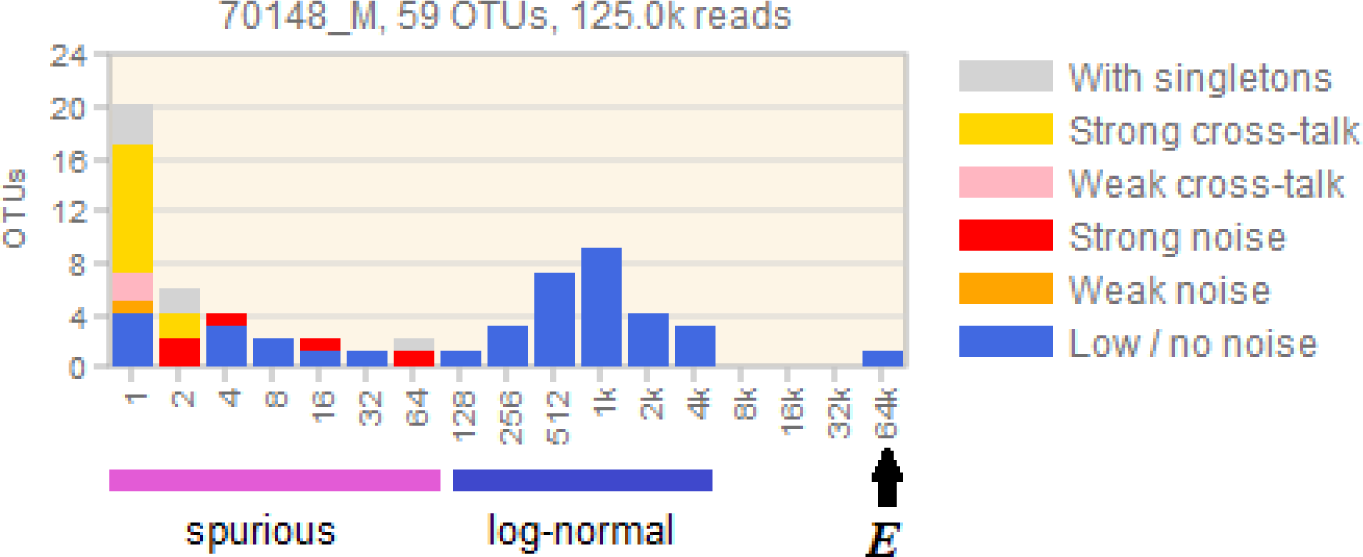
Octave plot of a representative prostate cancer tissue sample. This plot was generated from UPARSE OTUs of reads for sample 70148_M from (Yow *et al*., 2017). We interpret bins 1 to 64 as predominantly spurious and bins 128 through 4k as approximately log-normal. The log-normal shape appears to be robust because there is no evidence of spurious OTUs due to noise or cross-talk, and there is no excess when singletons are included (gray bars). The highly abundant *Escherichia/Shigella* OTU is indicated by ***E***; this OTU is found in all samples. We found no core microbiome in the log-normal range, as explained in the main text. Note that this plot covers a range of abundances of five orders of magnitude. The putatively spurious bins 1 through 64 contain 30 of the 59 OTUs, but only 254 of the 125,000 reads (0.2%).

### Octave plots of open-reference OTUs

Fig. 5 shows octave plots for OTUs generated by QIIME open-reference clustering for comparison with plots made with UPARSE OTUs from the same reads (Fig. 3). While the UPARSE histograms have distinctly different shapes (types *C*, *T*, *J* and *A*), the open-reference histograms have similar shapes (all type *J*). This is almost certainly explained by spurious open-reference OTUs which are increasingly prevalent at lower abundances and proliferate rapidly with increasing numbers of reads, causing an underlying true distribution of any shape to be overlaid by a larger *J*-shaped histogram containing the spurious OTUs (Fig. 6). The spurious overlay increases indefinitely as more reads are added, corresponding to a rarefaction curve which converges on a constant upwards gradient (the rate of spurious OTU generation) rather than a horizontal asymptote, as seen on the mock community tests in (Edgar, 2017b).

**Fig. 5.**
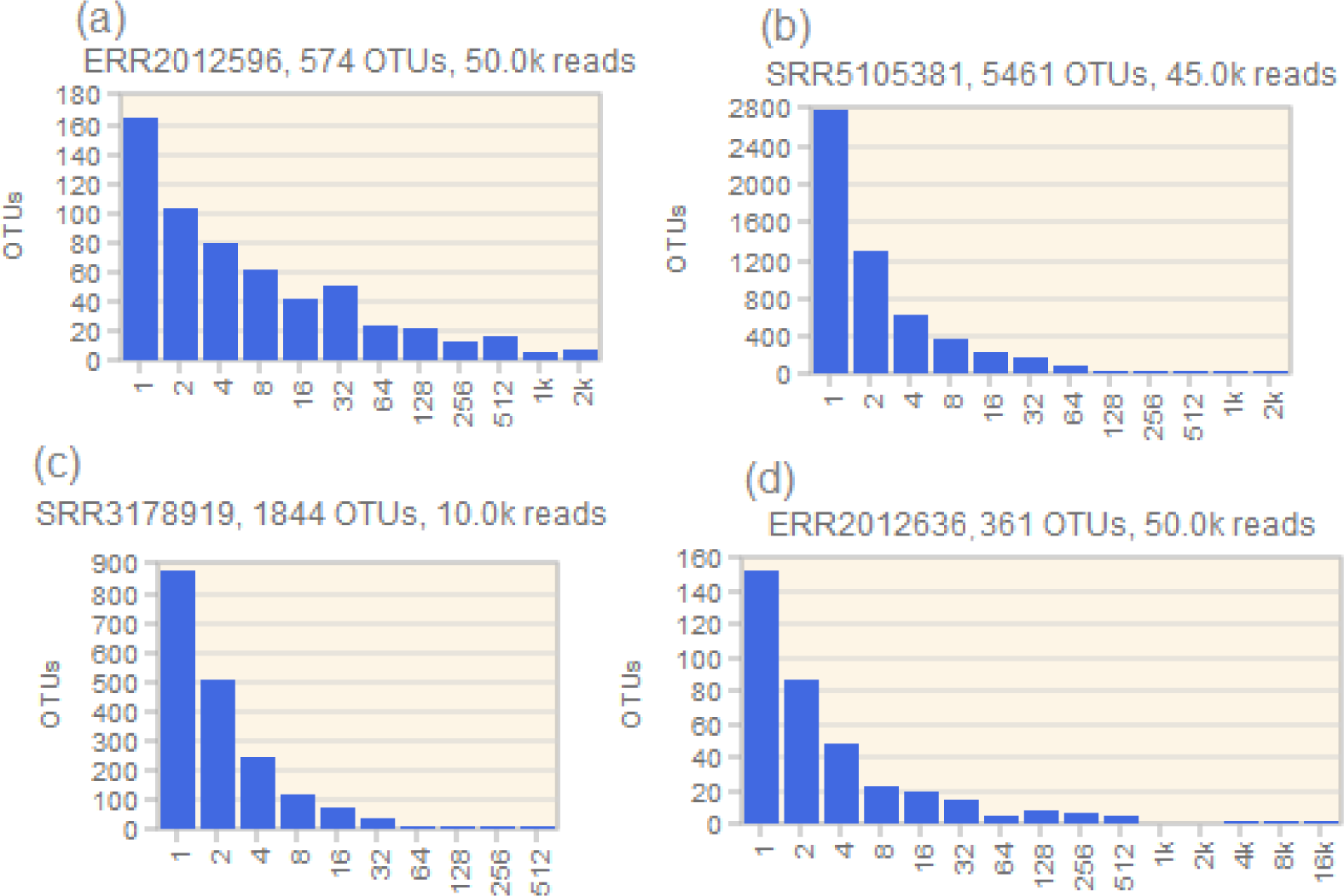
Octave plots of QIIME v1.9 open-reference OTUs for the same samples as Fig. 3. With open-reference OTUs, reported diversity is much greater and all distributions are *J*-shaped. This is almost certainly due to spurious OTUs, which are increasingly prevalent at low abundances.

**Fig. 6.**
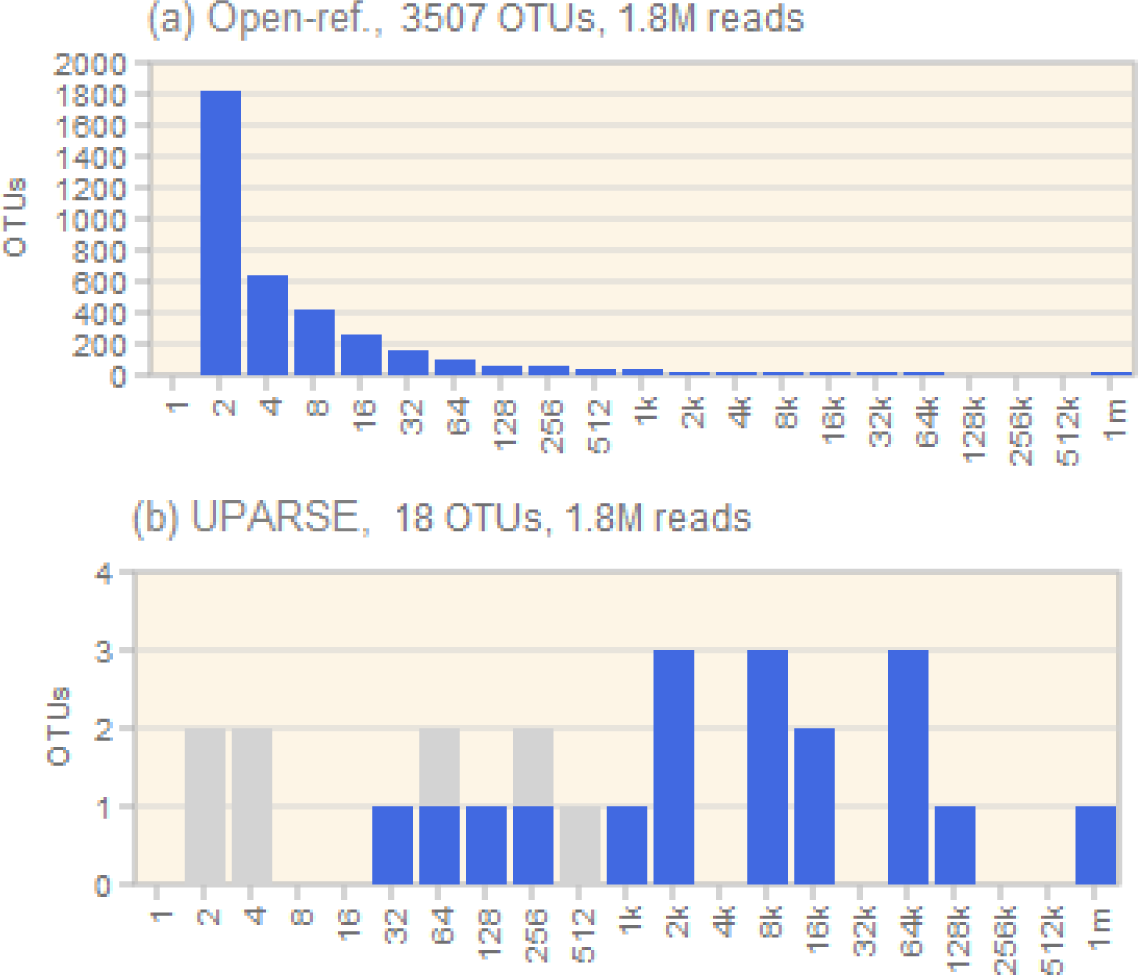
Octave plots for a mock community sample. These plots were generated from (a) QIIME v1.9 open-reference OTUs and (b) UPARSE OTUs from reads of the Staggered1 sample, which contains 21 strains. The first bin of the open-reference plot is empty because OTUs with exactly one read per dataset are discarded by open-reference clustering, and only one sample was included in the analysis. Both plots contain similar numbers of reads (1,750,658 for open-reference, 1,753,266 for UPARSE from a total of 1,857,075 reads in the FASTQ file), which UPARSE assigns to 18 OTUs compared to 3,507 OTUs for open-reference. With UPARSE, the singleton excess is seven additional OTUs (gray bars). With singletons included, the open-reference bin 1 contains 12,692 OTUs, making a total of 16,199 (not shown). These results show that the open-reference distributions seen in Fig. 5 can be explained by a large *J* shape due to spurious OTUs overlaid onto smaller true distributions. The UPARSE distribution is type *A* (anomalous), reflecting that the sample is not a natural interacting ecosystem. The UPARSE plot correctly indicates that the full diversity has been sampled because the first five bins are empty when singleton read sequences are excluded, while the open-reference plot misleadingly indicates that the true richness is much greater than the 3,507 reported OTUs.

## Discussion

### Extrapolating to unseen diversity

With shape *C*, a complete bell curve is seen, implying that most or all of the diversity has been enumerated. With shape *T*, a peak is apparent, implying that more than half the total richness has been enumerated (assuming an approximately symmetrical distribution). The downslope to the left of the peak indicates the shape of the low-abundance tail, and it is then reasonable to extrapolate to unseen OTUs, for example by fitting log-normal parameters (Bulmer, 1974). The Chao1 (Chao, 1984) non-parametric estimator is also reasonable providing that the numbers of singleton and doublet OTUs required by its formula can be accurately measured, but this is usually not possible in practice because low-abundance OTUs, especially singletons, are often spurious. With shape *J*, extrapolation is not supportable because distributions with a wide range of different forms and parameters can be fitted to the observed abundances, as illustrated by panels (c) and (d) of Fig. 1 which have similar shapes but are generated by log-normal and log-series respectively. Even if the distribution is assumed to be log-normal, it is not clear whether the peak is in bin 1 or in an unseen bin to the left of the *y* axis. With anomalous shapes, it is self-evident that extrapolation based on general principles, e.g. fitting to log-normal or calculating Chao1, is not supported by the data.

### Meaningful alpha diversity metrics

With an incomplete enumeration, the value of an alpha diversity metric calculated from observed OTUs may be substantially different from its true value, i.e. the value that would be obtained from a complete, unbiased, and error-free set of abundances. In practice, most enumerations appear to be incomplete, and the effects of errors and biases are difficult to determine. These issues are particularly acute with richness which may be arbitrarily under- or over-estimated due to OTUs that are missing (because of incomplete enumeration, failure to amplify due to primer mismatches, and discarding of low-abundance reads and/or low-abundance OTUs by the analysis pipeline) and OTUs that are spurious (due to unfiltered experimental errors such as bad sequences, chimeras, cross-talk, and contaminants). To a first approximation, it is reasonable to assume that experimental errors occur with comparable rates in samples which have been sequenced by the same protocol. If enumerations are complete, then given this assumption it is valid to compare observed richness between samples because the excess or deficit due to experimental error is approximately constant. However, regardless of errors, when enumerations are incomplete, a change in observed richness could be due to a change in AD shape and therefore does not necessarily imply a change in true richness. With the Chao1 estimator, issues with richness are compounded by the difficulties in accurately estimating the abundances of singleton and doublet OTUs. Evenness metrics such as Simpson’s index (Simpson, 1949) are suspect because they are sensitive to the presence of a dominant OTU which may have exaggerated abundance, e.g. due to high gene copy count. Changes in observed richness and/or shape are reflected by changes in Shannon entropy (Shannon, 1948), which down-weights OTUs with low frequencies and is therefore less sensitive to excesses or deficits at low abundances where experimental error is most likely to cause problems. Entropy is therefore our preferred metric. Numerical values of entropy and changes in entropy are opaque (i.e., have no immediate biological interpretation), but we regard this as an advantage because metrics which have more transparent interpretations in traditional biodiversity studies, such as richness or Simpson’s index, can be misleading when applied to metagenomic data. A statistically significant change in entropy implies a biologically significant change in diversity, which can be interpreted by reviewing AD shapes, e.g. in octave plots.

### Estimating read depth required to capture all OTUs

If, and only if, it is possible to extrapolate to unseen diversity, then it is also possible to estimate the number of reads required to obtain a complete enumeration, i.e. to capture all OTUs. With shape *C*, there is no evidence that additional reads are required. With *T*, the number of reads can be estimated from an octave plot by assuming that the distribution is symmetrical and determining the number of bins (*m*) that would appear to the left of the *y* axis (Fig. 7). The number of reads required to capture *m* additional bins is 2*^m^ N* where *N* is the number of reads used to generate the plot. With shapes *J* and *A*, there is insufficient data to make a reliable estimate.

**Fig. 7.**
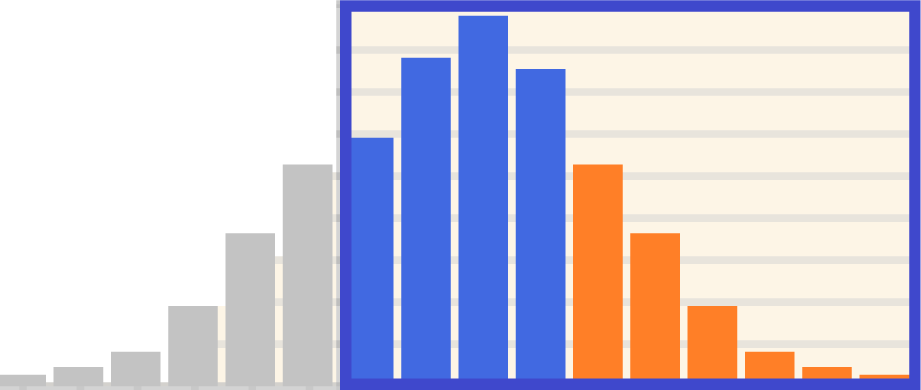
Estimating the number of additional reads required to capture all OTUs. The rectangle is an octave plot with truncated (*T*) shape. Assuming a symmetrical distribution, the tail on the right (orange) will be the same as the unseen tail to the left of the *y* axis (gray), giving an estimate of 6 unseen bins. This implies that a factor of 2^6^ = 64× more reads are required.

### Visualization enables new insights into microbial diversity

The term “octaves” for bins that double in size was introduced by Preston by analogy with musical notes which double in frequency in each successive octave (middle C is 262Hz, C’ is 524Hz, C” is 1048Hz and so on). The key modifications (pun intended) in our octave plots are the bin boundaries, which are critically important for maintaining the shape of an incompletely enumerated distribution, and coloring to indicate likely spurious OTUs.

Octave plots enable new insights into microbial abundance distributions, facilitating assessment of alpha diversity to suggest hypotheses and distinguish biologically meaningful inferences from artifacts of experimental error.

## References

Bokulich, N.A., Subramanian, S., Faith, J.J., Gevers, D., Gordon, J.I., Knight, R., Mills, D.A., and Caporaso, J.G. (2013). Quality-filtering vastly improves diversity estimates from Illumina amplicon sequencing. Nat. Methods 10, 57–9.

Bulmer, M.G. (1974). On Fitting the Poisson Lognormal Distribution to Species-Abundance Data. Biometrics 30, 101–110.

Callahan, B.J., McMurdie, P.J., Rosen, M.J., Han, A.W., Johnson, A.J., and Holmes, S.P. (2016). DADA2: High-resolution sample inference from Illumina amplicon data. Nat. Methods 13, 581.

Caporaso, J.G., Kuczynski, J., Stombaugh, J., Bittinger, K., Bushman, F.D., Costello, E.K., Fierer, N., Peña, A.G., Goodrich, J.K., Gordon, J.I., et al. (2010). QIIME allows analysis of high-throughput community sequencing data. Nat. Methods 7, 335–336.

Carini, P., Marsden, P.J., Leff, J.W., Morgan, E.E., Strickland, M.S., and Fierer, N. (2017). Relic DNA is abundant in soil and obscures estimates of soil microbial diversity. Nat. Microbiol. 2, 16242.

Carlsen, T., Aas, A.B., Lindner, D., Vraalstad, T., Schumacher, T., and Kauserud, H. (2012). Don’t make a mista(g)ke: Is tag switching an overlooked source of error in amplicon pyrosequencing studies? Fungal Ecology 5(6) 747–749.

Chao, A. (1984). Nonparametric estimation of the numbers of classes in a population. Scand. J. Stat. 11, 265–270.

Cho, I., and Blaser, M.J. (2012). The human microbiome: at the interface of health and disease. Nat. Rev. Genet. 13, 260–270.

Edgar, R.C. (2010). Search and clustering orders of magnitude faster than BLAST. Bioinformatics 26, 2460–1.

Edgar, R.C. (2013). UPARSE: highly accurate OTU sequences from microbial amplicon reads. Nat. Methods 10, 996–8.

Edgar, R.C. (2016). SINTAX: a simple non-Bayesian taxonomy classifier for 16S and ITS sequences. https://doi.org/10.1101/074161.

Edgar, R.C. (2017a). UNCROSS: Filtering of high-frequency cross-talk in 16S amplicon reads. Doi https://doi.org/10.1101/088666.

Edgar, R.C. (2017b). Accuracy of microbial community diversity estimated by closed- and open-reference OTUs. PeerJ 2017.

Edgar, R.C. (2017c). UNBIAS: An attempt to correct abundance bias in 16S sequencing, with limited success. https://doi.org/10.1101/124149.

Edgar, R.C. (2017d). UNOISE2: improved error-correction for Illumina 16S and ITS amplicon sequencing. https://doi.org/10.1101/081257.

Edgar, R.C. (2017e). Accuracy of microbial community diversity estimated by closed- and open-reference OTUs. PeerJ 5:e3889, https://doi.org/10.7717/peerj.3889.

Edgar, R.C. (2018). UNCROSS2, an improved algorithm for cross-talk detection and filtering.

Fisher, R.A., Corbet, A.S., and Williams, C.B. (1943). The relation between the number of species and the number of individuals in a random sample of an animal population. J. Anim. Ecol. 42–58.

Gray, J.S., Bjørgesæter, A., and Ugland, K.I. (2006). On Plotting Species Abundance Distributions. J. Anim. Ecol. 75, 752–756.

Hartmann, M., Niklaus, P.A., Zimmermann, S., Schmutz, S., Kremer, J., Abarenkov, K., Lüscher, P., Widmer, F., and Frey, B. (2014). Resistance and resilience of the forest soil microbiome to logging-associated compaction. ISME J. 8, 226–244.

Huse, S.M., Welch, D.M., Morrison, H.G., and Sogin, M.L. (2010). Ironing out the wrinkles in the rare biosphere through improved OTU clustering. Environ. Microbiol. 12, 1889–98.

Magurran, A.E. (2004). Measuring Biological Diversity. ISBN 978-0-632-05633-0.

May, R.M. (1975). Patterns of species abundance and diversity. In Ecology and Evolution of Communities, M.L. Cody, and J.M. Diamond, eds. (Cambridge, MA: Harvard University Press), pp. 81–120.

Moran, M.A. (2015). The global ocean microbiome. Science 347, aac8455.

Parulekar, N.N., Kolekar, P., Jenkins, A., Kleiven, S., Utkilen, H., Johansen, A., Sawant, S., Kulkarni-Kale, U., Kale, M., and Sæbø, M. (2017). Characterization of bacterial community associated with phytoplankton bloom in a eutrophic lake in South Norway using 16S rRNA gene amplicon sequence analysis. PLOS ONE 12, e0173408.

Pflughoeft, K.J., and Versalovic, J. (2012). Human microbiome in health and disease. Annu. Rev. Pathol. 7, 99–122.

Preston, F.W. (1948). The Commonness, And Rarity, of Species. Ecology 29(3) 254–283.

Rideout, J.R., He, Y., Navas-Molina, J. a, Walters, W. a, Ursell, L.K., Gibbons, S.M., Chase, J., McDonald, D., Gonzalez, A., Robbins-Pianka, A., et al. (2014). Subsampled open-reference clustering creates consistent, comprehensive OTU definitions and scales to billions of sequences. PeerJ 2, e545.

Shannon, C.E. (1948). A Mathematical Theory of Communication. Bell Syst. Tech. J. 27, 379–423.

Simpson, E.H. (1949). Measurement of diversity. Nature 163, 688.

Virtanen, S., Kalliala, I., Nieminen, P., and Salonen, A. (2017). Comparative analysis of vaginal microbiota sampling using 16S rRNA gene analysis. PLOS ONE 12, e0181477.

Whittaker, R.H. (1965). Dominance and diversity in land plant communities. Science 147(3655) 250–260.

Williamson, M., and Gaston, K.J. (2005). The lognormal distribution is not an appropriate null hypothesis for the species–abundance distribution. J. Anim. Ecol. 74, 409–422.

Yow, M.A., Tabrizi, S.N., Severi, G., Bolton, D.M., Pedersen, J., Giles, G.G., and Southey, M.C. (2017). Characterisation of microbial communities within aggressive prostate cancer tissues. Infect. Agent. Cancer 12, 4.

